# Dual functioning by the PhoR sensor is a key determinant to *Mycobacterium tuberculosis* virulence

**DOI:** 10.1101/2023.04.13.536687

**Authors:** Prabhat Ranjan Singh, Harsh Goar, Partha Paul, Khushboo Mehta, Bhanwar Bamniya, Anil Kumar Vijjamarri, Roohi Bansal, Hina Khan, Subramanian Karthikeyan, Dibyendu Sarkar

## Abstract

PhoP-PhoR empowers *M. tuberculosis* to adapt to diverse environmental conditions, and remains essential for virulence. Although PhoP and PhoR have been structurally characterized, the signal(s) that this TCS responds to remains unknown. In this study, we show that PhoR is a sensor of acidic pH/high salt conditions, which activate PhoP via phosphorylation. Transcriptomic studies uncover that acidic pH-inducible expression of PhoP regulon is significantly inhibited in a PhoR-deleted *M. tuberculosis*. Using genome-wide screening we further identify a non-canonical mechanism of PhoP phosphorylation by the sensor kinase PrrB. To investigate how phosphorylation of PhoP is regulated, we discovered that PhoR functions as a phosphatase. Our results identify the motif/residues responsible for contrasting kinase/phosphatase dual functioning of PhoP, and collectively determine the homeostatic regulation of intra-mycobacterial P~PhoP which controls the final output of PhoP regulon. Together, these data uncover that PhoR plays a central role in mycobacterial adaptation to low pH conditions within the host macrophage phagosome. Consistent with these results a PhoR-deleted *M. tuberculosis* remains significantly attenuated in macrophages and animal models.

## Introduction

*M. tuberculosis* is highly adaptable to complex and varying host environment it must encounter during infection. This adaptability largely relies on signal transducing systems which turn on complex transcription networks (Bretl *et al*, 2011). Bacterial two component systems (TCSs) are coupled signal sensing and transduction apparatus which comprise of two proteins, a transmembrane sensor kinase (SK) responsible for sensing signal(s) and a cognate cytoplasmic response regulator (RR), that translate environmental information into a physiological response (Capra & Laub, 2012; Casino *et al*, 2009). Underlying the functional diversity of TCSs is a common set of chemical reactions by which the activated RR impacts expression of numerous genes in response to signal(s), that is first sensed by a SK which then activates the cognate regulator via a sequential phosphotransfer reaction (Gotoh *et al*, 2010). *M. tuberculosis* genome encodes for 12 complete TCSs, fewer relative to other bacterial species with comparable genome size (Cole *et al*, 1998; Parish, 2014; Tucker *et al*, 2007).

Numerous studies have uncovered that several TCSs, namely DosRST, PhoPR, MprAB, SenX3-RegX3, PdtaRS and MtrAB, have defined roles to *in vivo* virulence, as determined by various infection models utilizing immune-compromised and competent mice, guinea pigs, rabbits and non-human primates (Banerjee *et al*, 2019; Buglino *et al*, 2021; Converse *et al*, 2009; Mehra *et al*, 2015; Parish *et al*, 2003; Rifat *et al*, 2014; Tischler *et al*, 2013; Walters *et al*, 2006). Of these, the PhoP/PhoR pair of proteins of *M. tuberculosis* form a specific TCS that functions with a key regulatory role in controlling cell-wall lipid composition and virulence (Gonzalo-Asensio *et al*, 2008b; Gonzalo Asensio *et al*, 2006; Ludwiczak *et al*, 2002; Perez *et al*, 2001), immediate and enduring hypoxic responses, aerobic and anaerobic respiration (Gonzalo-Asensio *et al*, 2008a; Singh *et al*, 2020a), secretion of major virulence factors (Anil Kumar *et al*, 2016; Frigui *et al*, 2008), stress response and persistence (Baker *et al*, 2014; Gonzalo-Asensio *et al*., 2008a; Sevalkar *et al*, 2019; Singh *et al*., 2020a) [for reviews see (Ryndak *et al*, 2008), (Broset *et al*, 2015)]. Notably, PhoP controls synthesis of three classes of polyketide-derived acyltrehaloses, namely sulfolipids, di-acyl trehaloses (DATs) and poly acyltrehaloses (PATs) (Gonzalo Asensio *et al*., 2006; Walters *et al*., 2006). In keeping with this, avirulent *M. tuberculosis* H37Ra with a genomic copy of PhoP harbouring a single nucleotide polymorphism (SNP) within its effector domain, shows growth attenuation (Frigui *et al*., 2008; Lee *et al*, 2008) and lacks polyketide-derived acyltrehaloses relative to virulent *M. tuberculosis* H37Rv (Chesne-Seck *et al*, 2008). More importantly, virulence and PAT/DAT biosynthesis is restored in *M. tuberculosis* H37Ra expressing copy of *M. tuberculosis* H37Rv *phoP* gene (Frigui *et al*., 2008; Goyal *et al*, 2011). Transcriptomic study reveals that ~2% of the *M. tuberculosis* H37Rv genome is regulated by PhoP at the transcriptional level (Solans *et al*, 2014; Walters *et al*., 2006). Therefore, with inactivation of *phoPR* the tubercle bacilli demonstrate a strikingly reduced multiplication in macrophages, and consequently the *phoPR* deletion strain is considered in vaccine trials (Arbues *et al*, 2013).

Studies aimed to understand functioning of the transcription factor uncovered that *ΔphoP*-H37Rv complemented with the phosphorylation defective PhoP fails to rescue complex lipid biosynthesis (Goyal *et al*., 2011), suggesting P~PhoP dependent regulation of mycobacterial genes (Goyal *et al*., 2011). However, we still do not fully understand the mechanism of activation of the virulence-associated PhoP/PhoR system. Although a great deal is known about the mechanism of functioning of PhoP, relatively much less is known about PhoR, the HK sensor of this complex. PhoR is a homodimer, and each subunit consists of two transmembrane helices which flank an N-terminal sensor domain (extra-cytosolic), a HAMP domain, a DHp (dimerization and phosphotransfer) domain, and a cytoplasmic CA (catalytic and ATP-binding) domain, respectively. The DHp domain plays a central role of the HK sensor function. These include interactions with the ATP binding domain and autophosphorylation at His 259, contacting the partner RR to transfer phosphate group (for the latter’s activation), and accepting signal(s) from the upstream sensor domain to regulate phosphorylation (Xing *et al*, 2017). While *in vitro* studies have shown autophosphorylation of PhoR and phospho-transfer to PhoP (Gupta *et al*, 2006), *in vivo* phosphorylation of PhoR and PhoP is yet to be demonstrated. More importantly, the signal(s) that activates PhoP/PhoR TCS remains unknown.

In this study, we show that PhoR is a direct sensor of acidic pH conditions, which subsequently activates PhoP via phosphorylation. Using mycobacterial protein fragment complementation (M-PFC)-based screening coupled with phosphorylation assays, we identify PrrB as a bona fide phospho-donor of PhoP. We also demonstrate that in addition to its kinase activity, PhoR functions as a phosphatase of P~PhoP, and this activity of PhoR connects signal-dependent pool of intra-mycobacterial P~PhoP with downstream functioning of the regulator. Using mutant PhoR proteins, we provide evidence to show that contrasting kinase and phosphatase functions of the PhoR remain critically important to determine the final output of the acidic pH-inducible PhoP regulon. Thus, PhoR functions remain critical for its central role in mycobacterial adaptation to low pH conditions within the host macrophage phagosome. In keeping with these results, a PhoR-deleted *M. tuberculosis* remains significantly attenuated in macrophages and animal models.

## Results

### Acidic pH and high salt conditions of growth promotes phosphorylation of PhoR and PhoP

Because SK of a TCS is often responsible for sensing environmental signal(s), abundance of the SK transcripts does not necessarily correlate with activation of a TCS. However, determining the amount of phosphorylated SK/RR represents a more direct measure of activation of the system (Choi & Groisman, 2017). Thus, to examine whether PhoR can sense activation signal(s) during *in vitro* growth conditions, we studied phosphorylation of PhoR using phos-tag gels (Barbieri & Stock, 2008). In this assay, phos-tag gels effectively resolve phosphorylated and unphosphorylated forms of the SK/RR in cell lysates of bacteria grown under specific environmental conditions. These experiments were carried out in WT-H37Rv background in which His-tagged PhoR was expressed ectopically from the promoter of the 19-kDa antigen of p19KPro (De Smet *et al*, 1999), and the amount of P~PhoR was detected by Western blotting using antibodies that recognize His-tagged PhoR (Fig. 1A). When WT bacilli was grown under acidic (pH 4.5) conditions, >90% of the total PhoR protein was in the phosphorylated (P~PhoR) form compared to ~5% of P~PhoR (compare lane 1 and lane 2) for cells grown under normal (pH 7.0) conditions. In sharp contrast, >90% of PhoR was available in the unphosphorylated form in mycobacterial cells grown in presence of 5 mM diamide (under oxidative stress) (lane 3). Further, P~PhoR accounted for ~90% of total PhoR in extracts of bacterial cells grown in presence of 250 mM NaCl (salt stress) (lane 4). From these results, we conclude that *M. tuberculosis* PhoR is capable of sensing low pH and high salt conditions of growth.

**Fig. 1:**
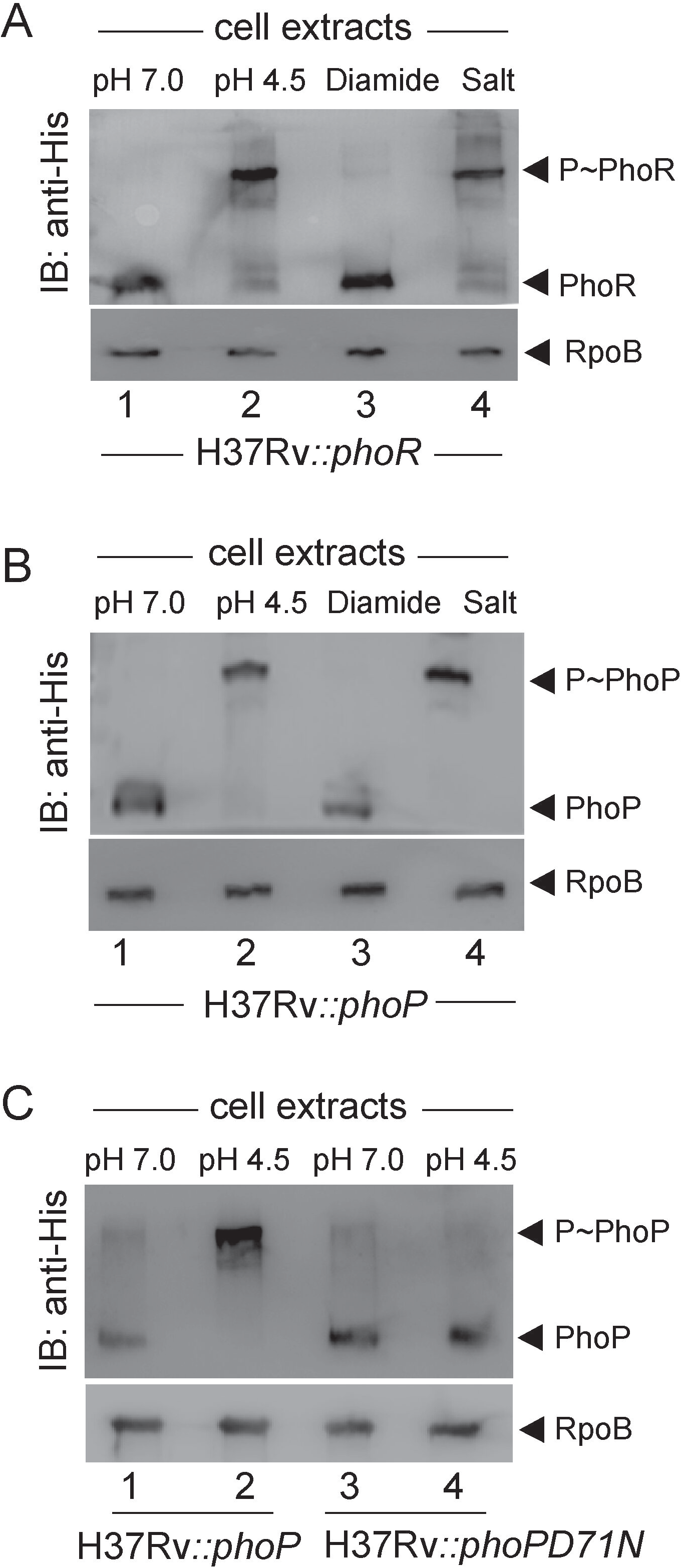
Mycobacterial growth under conditions of acidic pH and high salt concentration promote phosphorylation of PhoR and PhoP. Phos-tag analysis of cell lysates of WT-H37Rv harbouring His-tagged (A) PhoR, (B) PhoP or (C) phosphorylation-defective PhoPD71N proteins, were grown either under normal conditions (pH 7.0) or under indicated stress conditions, as described in the Methods. PhoR and PhoP were detected by Western blotting using anti-His antibodies, and RpoB (as a loading control) was detected in cell lysates using anti-RpoB antibody (Abcam). Data are representative of two independent experiments.

In order to probe phosphorylation of the cognate RR PhoP, we next expressed full-length *M. tuberculosis* PhoP protein in WT-H37Rv as described above. Cells were grown under stress conditions and presence of P~PhoP was probed in cell extracts using anti-His antibody (Fig. 1B). Remarkably, we could detect majority of P~PhoP (>80%) in extracts of mycobacterial cells grown under low pH (lane 2) or high salt (lane 4) conditions. In contrast, for cells grown under normal conditions (lane 1) or oxidative stress (lane 3), majority of the PhoP protein was in the unphosphorylated state. Thus, we conclude that stress-specific *in vivo* phosphorylation of PhoP is most likely dependent on phosphorylation of the cognate kinase, PhoR. As a control, we repeated an identical experiment using WT-H37Rv ectopically expressing His-tagged PhoPD71N, a phosphorylation defective mutant of PhoP (Fig. 1C). Although mycobacterial cells expressing WT-PhoP grown at pH 4.5 showed a significantly higher abundance of P~PhoP compared to cells grown at pH 7.0 (compare lane 1 and lane 2), we could not detect P~PhoP from mycobacterial cells expressing PhoPD71N (compare lane 3 and lane 4) under identical conditions. From these results, we conclude that low pH or high salt concentration promotes *in vivo* phosphorylation of *M. tuberculosis* PhoP. It should be noted that these results are consistent with a previous study suggesting synergistic response of *M. tuberculosis* H37Rv to Cl^-^ and low pH conditions as linked environmental cues (Tan *et al*, 2013).

### PhoP phosphorylation under acidic pH or high salt conditions requires the presence of PhoR

To examine whether PhoR is necessary for acidic pH and high salt concentration dependent phosphorylation of PhoP. We next utilized ‘mycobacterial recombineering’ (van Kessel & Hatfull, 2007) to construct a PhoR-deleted *M. tuberculosis* H37Rv (*ΔphoR*-H37Rv). The mutant construction is schematically shown in Fig. 2A, and described in the Methods. To verify the construct, PCR amplification was carried out using a pair of *phoR*-specific internal primers (FPphoRInt / RPphoRInt) and genomic DNA of *ΔphoR*-H37Rv (Fig. S1A). The results showed that *phoR*-specific amplicon was absent in the mutant relative to the WT-H37Rv (compare lane1 and lane 2). However, the complemented mutant utilizing an integrative plasmid harbouring a copy of *phoR* gene showed the presence of *phoR*-specific amplicon (lane 3). As controls, genomic DNA of these three strains yielded full-length *phoP*-specific amplicon (lanes 5-7). Also, *ΔphoR*-H37Rv was confirmed by Southern hybridization experiment (Fig. S1B). Although a *hrcA*-specific amplicon (~9.6-kb) could be detected in both WT and mutant strain using gene-specific end-labelled primers, *phoR*-specific (~2-kb) amplicon was detectable only in case of WT-H37Rv (compare lane 2 and lane 3). Consistent with these results, RT-qPCR experiments show that mRNA level of *phoR* remains undetectable in the mutant relative to WT-H37Rv, and expression of PhoR is significantly restored in the complemented mutant (Fig. 2B). The oligonucleotides used in cloning/amplifications, and the plasmid constructs used in expression of fusion proteins are listed in Tables S1, and S2, respectively. Next, *in vivo* phosphorylation assays using *ΔphoR*-H37Rv under varying conditions of growth showed that P~PhoP was undetectable in low pH or high salt conditions of growth (lanes 1-4, Fig. 2C), suggesting significantly reduced phosphorylation of PhoP in a PhoR-deleted strain, However, under identical conditions we could detect effective phosphorylation of PhoP in WT-H37Rv grown under low pH conditions (compare lanes 5 and 6). From these results, we conclude that abundance of PhoP~P during mycobacterial growth under acidic pH depends on the presence of the cognate kinase PhoR.

**Fig. 2:**
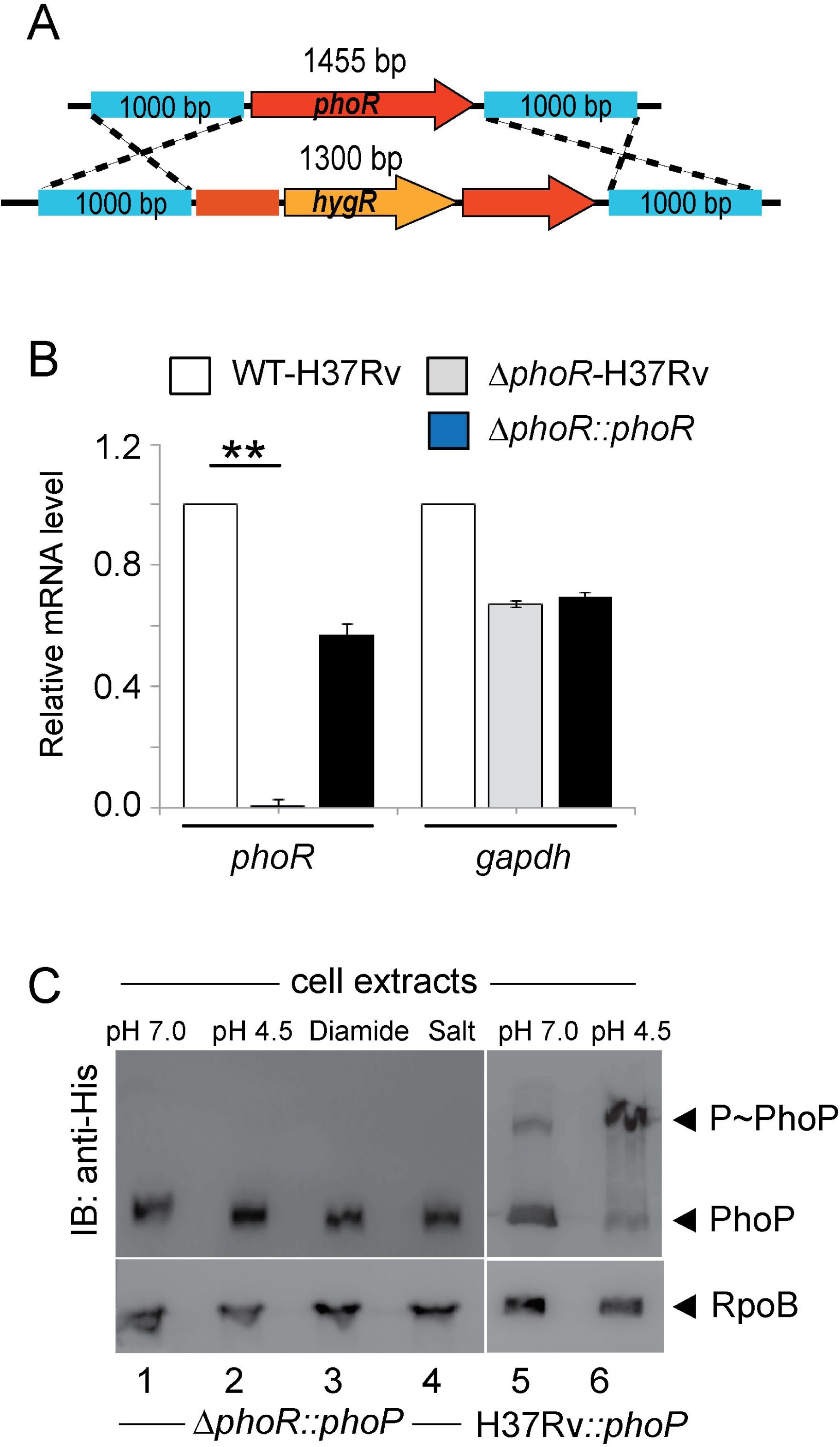
Acidic pH/high salt concentration-dependent phosphorylation of PhoP requires the presence of PhoR. (A) Schematic diagram showing construction of PhoR-deleted *M. tuberculosis* H37Rv (*ΔphoR*-H37Rv) by ‘mycobacterial recombineering’. (B) We utilized RT-qPCR experiments to compare mRNA level of *phoR* in the mutant and WT-H37Rv, as described in the Methods. Importantly, expression of *phoR* remains undetectable in the mutant relative to WT-H37Rv, but is significantly restored in the complemented mutant. As a control, *gapdh* levels remain comparable in WT-H37Rv, *ΔphoR*-H37Rv mutant and the complemented mutant strain. The results are derived from average of biological duplicates, each with one technical repeat (**P≤0.01). (C) To probe *in vivo* phosphorylation, Phos-tag Western blot analysis were performed using cell lysates prepared from *ΔphoR*-H37Rv, and WT-H37Rv harbouring His-tagged PhoP protein, respectively. The relevant strains were grown under normal or indicated stress conditions, as described in the Methods. Sample detection and data analysis were carried out as mentioned in the legend to Fig. 1.

### PhoR impacts global expression of the PhoP regulon

Numerous studies have shown that *M. tuberculosis* growth and transcriptional control are tightly regulated by environmental pH (Abramovitch *et al*, 2011; Vandal *et al*, 2009; Vandal *et al*, 2008). Along the line, global transcriptional response to acid stress suggests possible involvement of multiple regulators including *phoPR* in response to the cue of phagosomal acidification. (Bansal *et al*, 2017; Gonzalo Asensio *et al*., 2006; Rohde *et al*, 2007; Walters *et al*., 2006). To investigate regulatory influence of PhoR on mycobacterial gene expression, we compared transcriptomes of WT- and *ΔphoR*-H37Rv, grown under normal (pH 7.0) and acidic (pH 4.5) conditions of growth. The RNA-sequencing results (see Tables S3-S6) demonstrate that ~167 genes of WT-H37Rv are induced under low pH (pH 4.5) relative to mycobacterial cells grown under normal pH (pH 7.0) (Figs. S2A-B; also see Tables S3-S4). However, under identical conditions, *ΔphoR*-H37Rv displayed a low pH-inducible expression of only ~ 35 genes (Fig. 3A), suggesting that expression of ~79% of acidic pH-inducible genes require mycobacterial *phoR* locus (see Tables S4 and S6). Figs. S2C-D shows the Volcano plots of RNA-seq data assessing pH inducible expression of genes for WT-H37Rv and *ΔphoR*-H37Rv, respectively. Importantly, majority of the PhoR-controlled genes including genes from diverse functional categories like regulatory proteins, components of information pathways, virulence, detoxification and adaptation, metabolism and respiration, cell wall biosynthesis, lipid metabolism, and PE/PPE proteins etc., were part of the PhoP regulon.

**Fig. 3:**
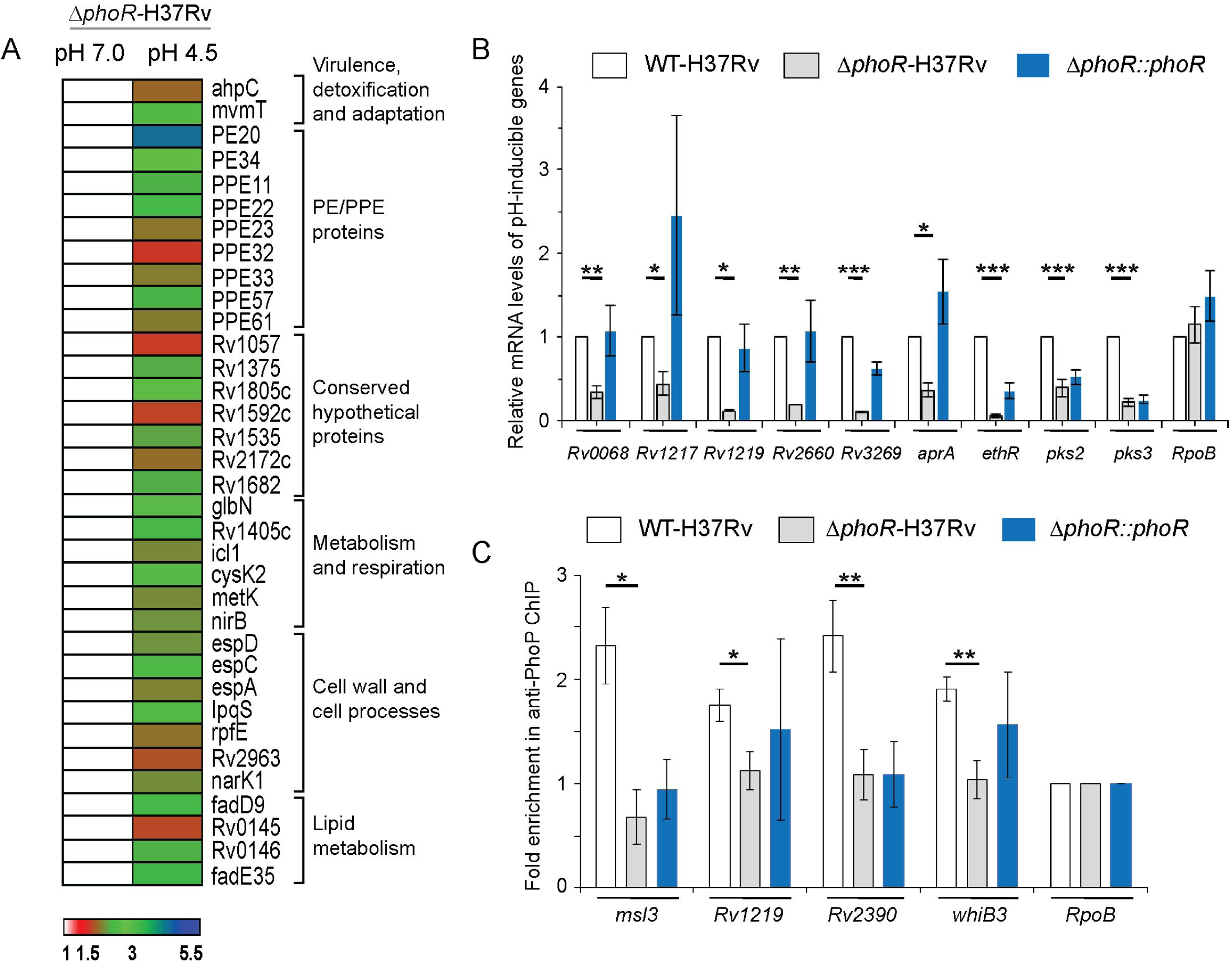
Global expression of acid-inducible mycobacterial genes require the *phoR* locus. (A) RNA-sequencing derived heat-map showing ~ 35 low pH-inducible genes that are differentially regulated in *ΔphoR*-H37Rv (>1.5 fold; p< 0.05) grown under low pH (pH 4.5) conditions compared to normal conditions (pH 7.0) of growth. The data, which represent average of two biological replicates, list significantly activated pH-inducible genes, majority of which are part of the PhoP regulon. (B) PhoR-dependent expression of acidic pH-inducible genes were examined by comparing expression of representative mycobacterial genes in WT-H37Rv, *ΔphoR*-H37Rv, and complemented strain, grown under normal and acidic pH conditions. The data show plots of average values from biological duplicates, each performed with one technical repeat (*P≤0.05; **P≤0.01; ***P≤0.001). (C) To examine *in vivo* recruitment of PhoP within its target promoters, ChIP was carried out using anti-PhoP antibody followed by qPCR using IP samples from WT-H37Rv and *ΔphoR*-H37Rv (see Methods section for further details). To assess fold enrichment, each data point was compared with the corresponding IP sample without adding antibody. The experiments were carried out as biological duplicates, each with at least one technical repeat (*P≤0.05; **P≤0.01). Non-significant differences are not indicated.

Next, we measured mRNA levels of representative low pH-inducible genes in *ΔphoR*-H37Rv relative to WT-H37Rv (Fig. 3B). In agreement with the RNA-seq data, RT-qPCR experiments demonstrate a significant down-regulation of representative PhoP regulon genes in *ΔphoR*-H37Rv. The fact that PhoR functions as an activator of these genes was further confirmed as stable expression of *phoR* in the complemented mutant could largely restore low pH-inducible gene expression. To assess whether presence of the *phoR* locus impacts on DNA binding activity of PhoP, we compared recruitment of the regulator *in vivo* within its promoters in WT-H37Rv and *ΔphoR*-H37Rv by chromatin immunoprecipitation (ChIP) assay (Fig. 3C). While qPCR measurements showed effective recruitment of PhoP in WT-H37Rv, under identical conditions a significantly weaker PhoP recruitment to its target sites was apparent in *ΔphoR*-H37Rv. Thus, we conclude that *in vivo* recruitment of PhoP requires the cognate kinase, PhoR. These results are consistent with previously-reported findings suggesting (a) PhoR as an obligate SK of PhoP (Xing *et al*., 2017) and (b) importance of phosphorylation of PhoP in high-affinity DNA binding and transcription regulation (Goyal *et al*., 2011). The sequences of oligonucleotides utilized in RT-qPCR and ChIP experiments are listed in Table S7.

### Screening for PhoP-interacting mycobacterial SK(s)

Cross-talk between TCS proteins has been considered a potential mechanism for complex stimulus integration and signal transduction in bacteria because of considerable high sequence and structural similarity in members of SK and RR family of proteins (Agrawal *et al*, 2015; Lee *et al*, 2012; Malhotra *et al*, 2015). Thus, we hypothesized additional non-canonical interactions and possible phospho-transfer between non-cognate SK(s) and PhoP. To probe SK-PhoP interaction(s), we utilized mycobacterial protein fragment complementation (M-PFC) using *M. smegmatis* as the surrogate host (Singh *et al*, 2006) (Figs. 4A-B). According to this assay, two interacting mycobacterial proteins which are expressed as C-terminal fusions with complementary fragments of mDHFR (murine dihydrofolate reductase), provides bacterial resistance to trimethoprim (TRIM) because of functional reconstitution of the full-length enzyme. *M. smegmatis* harbouring the corresponding plasmids were grown on 7H10/Kan/Hyg in the presence or absence of 15 µg/ml TRIM. Strikingly, *M. smegmatis* cells expressing both PrrB/PhoP proteins displayed clear growth in presence of TRIM, whereas none of the other *M. tuberculosis* SKs under identical conditions, displayed *in vivo* interaction with PhoP. These results suggest specific protein-protein interaction between PrrB and PhoP. Importantly, *M. smegmatis* harbouring vectors lacking an insert (empty) did not grow on 7H10/TRIM plates, while all other *M. smegmatis* strains displayed normal growth on 7H10 plates lacking TRIM. Tables S8 and S9 list the sequences of oligonucleotides, and relevant plasmids, respectively, used in cloning for M-PFC assays.

**Fig. 4:**
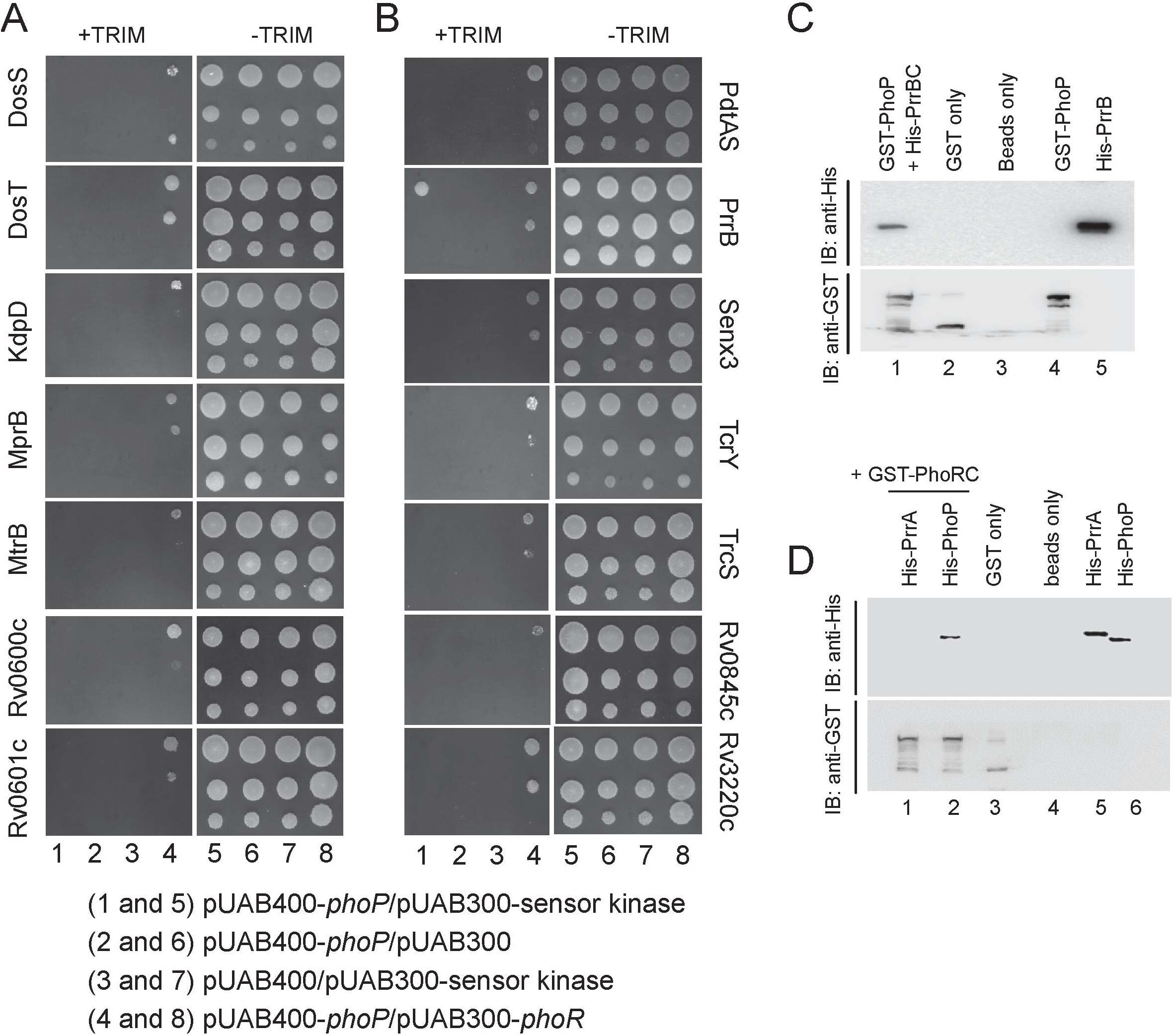
Screening to probe PhoP interacting mycobacterial SK(s). (A-B) M-PFC experiments co-expressing *M. tuberculosis* PhoP and SKs were used as a screen using *M. smegmatis* as the surrogate host. Co-expression of pUAB400-*phoP*/pUAB300-SK (indicated) pair (columns 1 and 5) relative to empty vector controls, pUAB400-*phoP*/pUAb300 (columns 2 and 6), or pUAB400/pUAB300-SK (columns 3 and 7), supporting *M. smegmatis* growth in presence of TRIM is suggestive of specific interaction. While co-expression of pUAB400-*phoP*/pAUB300-*phoR* (columns 4 and 8) in *M. smegmatis* displaying growth in presence of TRIM served as a positive control, growth of all the strains in absence of TRIM (columns 5-8) validated the assay. (C-D) To examine interactions *in vitro*, purified His_6_-PrrB or purified His_6_-PrrA and His_6_-PhoP were incubated with glutathione-Sepharose that was bound to GST-PhoP (C) or GST-PhoRC (D), respectively. Bound proteins (lane 1, panel C or lanes 1-2, panel D) were detected by Western blot using anti-His (upper panel) or anti-GST (lower panel) antibodies. Replicate experiments used glutathione Sepharose bound to GST (lane 2, panel C and lane 3, panel D), or the resin alone (lane 3, panel C or lane 4, panel D); lanes 4 and 5, panel C resolve GST-PhoP and His-PrrB, respectively; lanes 5, and 6, panel D resolve His-PrrA, and His-PhoP, respectively.

We also verified PrrB-PhoP interaction *in vitro* by pull-down assays using GST-PhoP and His_6_-tagged PrrBC (cytoplasmic region of PrrB comprising C-terminal residues 200 to 446) (Fig. 4C), cloned, expressed and purified as described in the Materials and Methods. In this experiment, GST-tagged PhoP was bound to glutathione-Sepharose, and then incubated with purified PrrBC. Upon elution of column-bound proteins, both proteins were detected in the same fraction (lane 1). However, a detectable signal was absent in case of only GST-tag (lane 2) or only resin (lane 3), further suggesting that PhoP interacts with PrrB. Also, we have carried out a pull-down experiment using GST-tagged PhoRC and His-tagged PrrA (Fig. 4D). Here, we were unable to detect presence of His-PrrA and GST-PhoRC in the same fraction (lane 1) although GST-PhoRC and His-PhoP coeluted in the same fraction (lane 2). From these results we conclude that although PhoR does not appear to interact with PrrA, *M. tuberculosis* PrrB interacts with PhoP.

### Phosphorylation of PhoP by a non-cognate SK

Having shown a specific interaction between PrrB and PhoP by M-PFC and *in vitro* pull-down assays, we investigated possible phosphotransfer between the two non-cognate partners. During *in vitro* experiments, PhoRC and PrrBC comprising C-terminal 193-485 and 200-446 residues of PhoR and PrrB, respectively, were used as His_6_-tagged recombinant SKs, whereas PhoP N-terminal domain (PhoPN, residues 1-141) and PrrA (residues 1-236) (see Table S2) were used as cognate RRs. Note that PhoPN was used in phosphotransfer assays because of comparable size of PhoP and PrrBC constructs (~247 amino acids). Auto-phosphorylation of PhoRC/PrrBC and subsequent phospho-transfer to PhoPN/PrrA were performed as described previously (Gupta *et al*., 2006; Pathak *et al*, 2010) and are schematically shown in Fig. 5A. For autophosphorylation, ~ 2.5 µM PhoRC/PrrBC was incubated in a phosphorylation mix containing ~25 µM γ^32^P-ATP. After incubation of 30 minutes at 25°C, ~ 5 µM purified PhoPN or PrrA was added to the phosphorylation mix, samples were incubated for indicated times, aliquots withdrawn, and the products analysed by SDS-PAGE. Importantly, recombinant P~PrrBC could effectively phosphorylate PhoPN (lanes 2-5, Fig. 5B). However, under identical conditions P~PhoRC was unable to phosphorylate recombinant PrrA (lanes 7-10, Fig. 5B). Notably, this observation is consistent with lack of detectable interaction between PhoRC and PrrA (Fig. 4D). Together, these results suggest specificity of PrrB-dependent phosphorylation of PhoP.

**Fig. 5:**
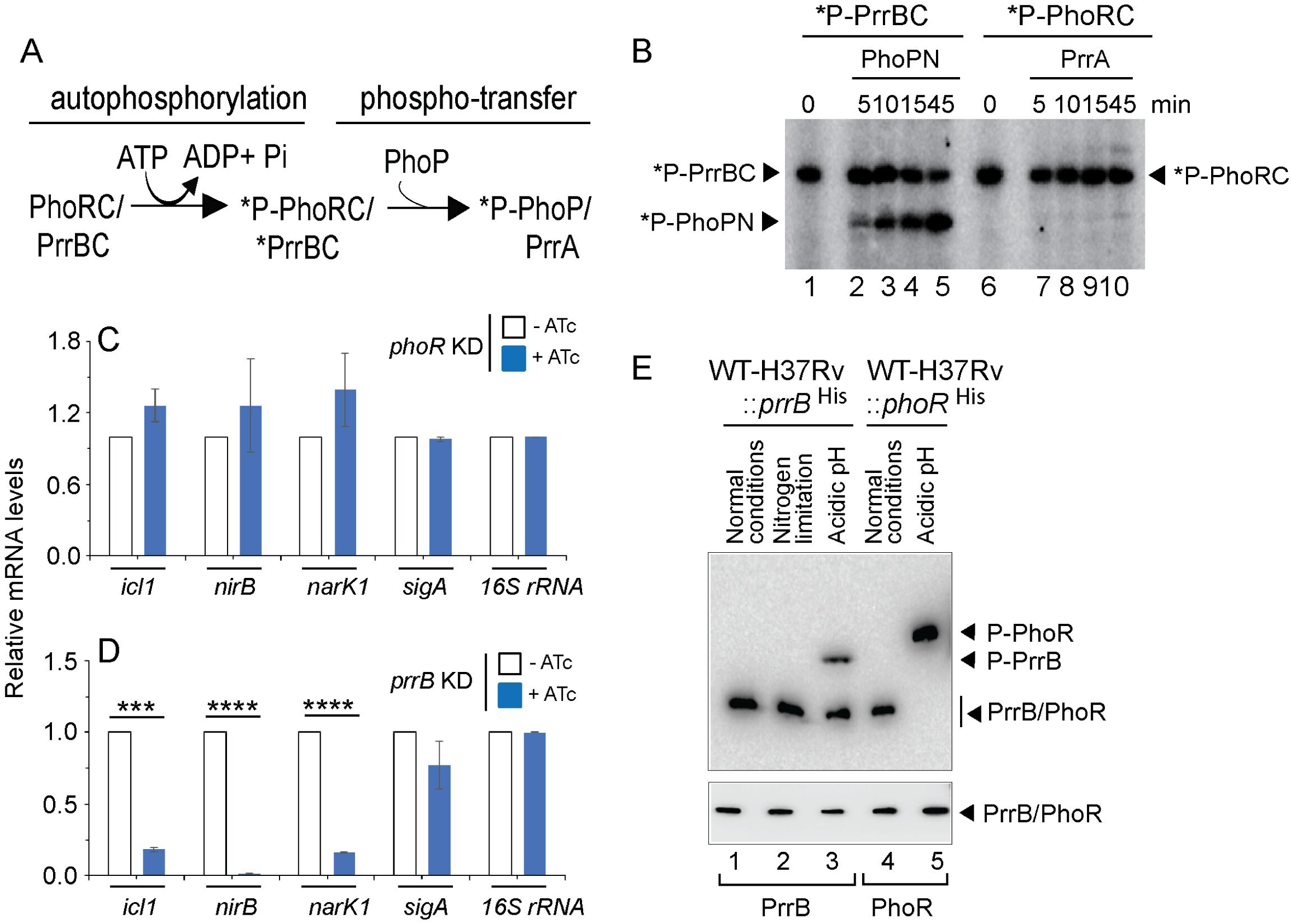
A non-canonical pathway of phosphorylation of PhoP. (A) Schematic diagram of auto-phosphorylation of SKs and subsequent phospho-transfer to RRs. (B) Phosphotransfer assays from radio-labelled PrrBC to PhoPN (lanes 1-5) and PhoRC to PrrA (lanes 6-10) are detailed in the Methods. Following SDS-PAGE analyses the reaction products were digitized by a phosphorimager (GE Healthcare); lane 1 and lane 6 resolve radio-labelled PrrBC and PhoRC, respectively. (C-D) mRNA levels of a few representative pH-inducible genes were next determined in the (C) *phoR* and (D) *prrB* knock-down constructs, respectively, grown under low pH conditions of growth. Each value is an average of duplicate measurements originating from biological duplicates (***P<0.001; ****P<0.0001). RT-qPCR measurements were performed as described in the Materials and Methods. (E) Phos-tag Western blot analysis of cell lysates prepared from WT-H37Rv harbouring His-tagged *prrB*. The bacterial strain was grown under normal conditions (pH 7.0) or indicated stress conditions, as described in the Methods, and the proteins of interest were detected by anti-His antibody. Data are representative of two independent experiments. As a loading control, His-PrrB or His-PhoR was detected from comparable amounts of cell lysates by Western blotting using anti-His antibody.

Next, to further probe activation of PhoP by specific SKs, we adopted a CRISPRi-based gene knock-down approach (Singh *et al*, 2016). The *phoR* and *prrB* knock-down constructs are described in the Methods, and relative mRNA levels compared expression of genes of interest in the corresponding knock-down strains relative to WT-H37Rv (Fig. S3A). We next grew these strains under low pH conditions to examine expression of a few representative genes, which showed *phoR* - independent pH-inducible activation (Fig. 5C, see Tables S3-S4). It should be noted that consistent with RNA-seq data from *ΔphoR*-H37Rv (Fig. 3A), *phoR* KD mutant fails to impact expression of these PhoP regulon genes (Fig. 5C). Fig. S3B shows RNA-seq data derived heat-map of these genes from WT- and *ΔphoR*-H37Rv, under low pH conditions of growth. However, in striking contrast, *prrB* KD shows a significant inhibition of expression of these genes relative to the WT bacilli under identical conditions (Fig. 5D). These results suggest that activation of PhoP regulon expression under acidic pH is partly attributable to mycobacterial PrrB. The above finding is consistent with PrrB - dependent phosphorylation of PhoP, and these results for the first time connect two SKs to integrate signal-dependent activation of PhoP, perhaps uncovering an effective means of mycobacterial adaptability to varying intracellular environments.

In an attempt to investigate under which condition PrrB is phosphorylated, we next grew WT-H37Rv expressing His-tagged PrrB under nitrogen limiting conditions and acidic pH, respectively, and assessed *in vivo* phosphorylation of the sensor kinases by phos-tag analysis (Fig. 5E). We included nitrogen limiting conditions, because mycobacterial PrrAB was reported to be induced under nitrogen limiting conditions (Haydel *et al*, 2012). Also, a recent study uncovered that PrrA as a transcription factor modulates mycobacterial stress response during low pH conditions (Giacalone *et al*, 2022). Expectedly, PrrB was mostly in the unphosphorylated form under normal conditions (lane 1) of growth. Also, we failed to observe phosphorylation of PrrB in cells grown under nitrogen limiting conditions (lane 2). However, PrrB was partially (~50%) phosphorylated under acidic pH relative to normal conditions of growth (compare lanes 4 and 5). This observation is remarkably consistent with PrrB-dependent regulation of acidic pH-inducible gene expression (Fig. 5D). In keeping with results shown in Fig. 1, as a control, PhoR undergoes robust phosphorylation under acidic pH conditions of growth (compare lanes 4 and 5, Fig. 5E). Together, these results uncover that acidic pH is also sensed by *M. tuberculosis* PrrB, which in turn phosphorylates PhoP.

### Probing dual functioning of sensor kinase PhoR

X-ray structure of *M. tuberculosis* PrrA, the closest family member of PhoP reveals that phosphorylation of the regulator appears to impact DNA binding and transcriptional control by the regulator (Nowak *et al*, 2006). Consistent with this, phosphorylation of PhoP strikingly impacts mycobacterial cell shape and morphology via its major regulatory control of genes responsible for complex lipid biosynthesis (Goyal *et al*., 2011). However, till date, it remains to be determined how cytoplasmic pool of phospho-PhoP is regulated. In the absence of an inducing stimulus, dephosphorylation of RR is critically important to ensure the TCS pathway gets reset. To examine the possibility that PhoR can function as a phosphatase, we designed an *in vitro* assay as shown schematically in Fig. 6A. In this experiment, PhoP was phosphorylated by radio-labelled ATP and acetyl-kinase as described elsewhere (Kaur *et al*, 2016), and incubated with equimolar PhoRC [comprising C-terminal 293 residues of PhoR (Gupta *et al*., 2006)] for varying lengths of time (Fig. 6B-E). Interestingly, we observed a time-dependent dephosphorylation of P~PhoP in presence of PhoRC (Fig. 6B). However, heat-denatured PhoRC, under identical conditions failed to dephosphorylate P~PhoP (Fig. 6C). As a control, recombinant PrrBC failed to dephosphorylate P~PhoP (Fig. 6D), suggesting PhoR-specific phosphatase activity. As an additional control, PhoRC was incubated with radio-labelled P~DosR under identical conditions (Fig. 6E). The results unambiguously demonstrate that PhoRC is incapable of dephosphorylating P~DosR. From these data, we surmise that PhoR functions as a specific phosphatase to maintain intra-mycobacterial cellular pool of P~PhoP.

**Fig. 6:**
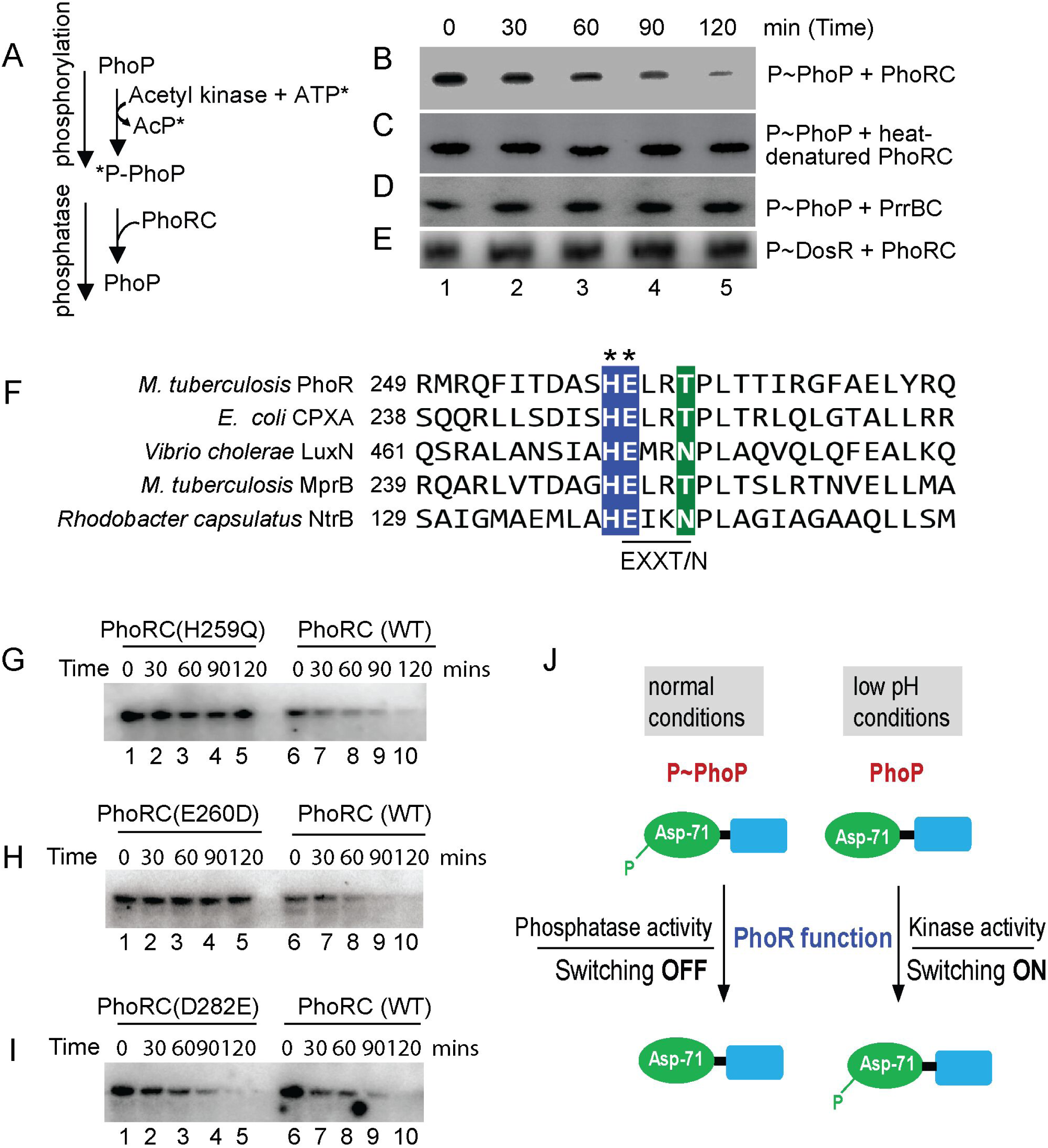
Identifying motif/residues responsible for phosphatase function of PhoR. (A) Scheme showing phosphorylation of PhoP by γ-^32^P-ATP and acetyl kinase and dephosphorylation of P-PhoP. (B-E) Time-dependent dephosphorylation of P-PhoP was examined by adding recombinant PhoRC (B), heat-denatured PhoRC (C), and purified PrrBC (D) as described in the Methods. (E) To examine specificity of newly-identified PhoR phosphatase activity, P~DosR was used as a substrate to examine possible dephosphorylation by adding purified PhoRC. The reactions were followed by SDS-PAGE analyses and digited by a phosphorimager. Each data is representative of at least two independent experiments. (F) DHp domain of PhoR was aligned with other SKs, which displayed phosphatase activity. The most conserved residues of the EXXT/N motif are indicated by asterisks. (G-I) To investigate the effect of mutations on the phosphatase function of PhoRC, purified WT and mutant proteins were incubated with P~PhoP for indicated times as described in Fig. 6B. In all cases, reactions were followed by SDS-PAGE analyses, and digitized by phosphorimager. (J) Schematic model showing balancing act of dual functioning of PhoR as a kinase (activation) and phosphatase (repression). Mutant PhoR protein, defective for kinase activity, fails to phosphorylate PhoP and thereby impact acid-inducible expression of the PhoP regulon. In contrast, phosphatase activity of PhoR dephosphorylates P~PhoP either to reverse active mycobacterial PhoP regulon in the absence of an inducing signal or to prevent unnecessary ‘triggering on’ of PhoP regulon. In summary, PhoR regulates net phosphorylation of PhoP by its dual functioning (kinase/phosphatase) to control context-sensitive expression of the PhoP regulon.

To determine which residue(s) of the DHp domain of PhoRC contribute to phosphatase activity, we next aligned α1-helix of the DHp domain of PhoR sequence with its family members, known to have exhibited phosphatase activity (Fig. 6F). A closer inspection of the sequence comparison reveals the presence of a conserved EXXT/N motif adjacent to His-259, the primary phosphorylation site of PhoR (Gupta *et al*., 2006). Remarkably, this is consistent with a few orthologues such as, EnvZ (Zhu *et al*, 2000) and CPXA (Raivio *et al*, 1999), which exhibit cognate RR-specific phosphatase activity attributable to either of the sequence motifs DXXXQ or E/DXXT/N, present within the α1 helix of DHp domain (Kaur *et al*., 2016; Willett & Kirby, 2012). Fig. S4A shows the structural model of PhoRC indicating the most conserved residues of α1 helix.

To examine role of the EXXT/N motif (residues 260-263) of PhoR in phosphatase activity, we replaced the most conserved residue (E260) by site-directed mutagenesis, and constructed a point mutant, PhoRCE260D. As a control, we also constructed PhoRCD282E, a mutant PhoRC replacing an aspartate with a glutamate further downstream of the α1 helix. The WT- and mutant PhoRC proteins were purified (Gupta *et al*., 2006), and *in vitro* autophosphorylation experiments using recombinant proteins demonstrate that except PhoRCH259Q (lane 1, Fig. S4B), the other two mutants (lanes 2-3) undergo auto-phosphorylation as that of WT PhoRC (lane 4). This is consistent with His-259 as the primary site of phosphorylation of PhoR (Gupta *et al*., 2006). Also, these results further suggest that the point mutants are structurally stable and functionally active.

Next, phosphatase assays were carried out as described in Figs. 6B-E using equimolar WT- or mutant PhoRC (recombinant) proteins (Fig. 6G-I). As expected, phosphorylation-defective PhoRCH259Q was unable to function as a phosphatase (lanes 1-5, Fig. 6G), whereas WT-PhoRC showed efficient phosphatase activity under identical conditions (lanes 6-10, Fig. 6G). Interestingly, PhoRCE260D was found to be remarkably ineffective for phosphatase activity relative to WT-PhoRC (compare lanes 1-5, and lanes 6-10, Fig. 6H). In contrast, PhoRCD282E, under identical conditions examined, displayed a comparable phosphatase activity as that of WT-PhoRC (compare lanes 1-5, with lanes 6-10, Fig. 6I). Thus, we conclude that although H259 remains the primary phosphorylation site, the adjacent conserved residue E260 of PhoR is critically required for phosphatase activity of the SK. Together, these results suggest importance of a limited number of residues of the DHp α1-helix in kinase/phosphatase dual functioning of PhoR. Based on these results, we present a schematic model (Fig. 6J) of pH-dependent mycobacterial adaptive program via phosphorylation-coupled ‘switching on’ and dephosphorylation-mediated ‘switching off’ of the PhoP regulon utilizing contrasting kinase and phosphatase activities of PhoR, respectively.

### PhoR is essential for mycobacterial survival in macrophages and mice

Having shown that PhoR is essential for phosphorylation of PhoP as well as maintenance of P~PhoP homeostasis in a pH-dependent mechanism, we sought to investigate whether PhoR contributes to mycobacterial virulence. Using a mice infection experiment, a previous study by Wang and co-workers had shown that PhoR-deleted *M. tuberculosis* displayed a lower lung burden of the bacilli relative to WT-H37Rv (Xing *et al*., 2017). We used WT-H37Rv and *ΔphoR*-H37Rv to infect murine macrophages (Fig. 7A). Although WT-H37Rv could effectively inhibit phagosome-lysosome fusion, *ΔphoR*-H37Rv readily matured into phagolysosomes, strongly indicating increased trafficking of the mycobacterial strain to lysosomes. To examine whether *ΔphoR*-H37Rv is growth defective or growth attenuated in macrophages, we compared survival of WT-H37Rv and *ΔphoR*-H37Rv within macrophages 3- and 48-hrs post infection (Fig. S5). Notably, *ΔphoR*-H37Rv shows a comparable growth 3-hr post infection as that of WT-H37Rv. However, 48 hr post infection the mutant shows a significantly reduced growth (~3.7-fold) relative to WT-H37Rv. More importantly, growth-attenuation is significantly rescued in the complemented mutant strain, suggesting that *phoR* locus is essential for mycobacterial growth in macrophages. In keeping with these results, bacterial co-localization data (Fig. 7B) and Pearson’s plot (Fig. 7C) strongly suggest the *phoR* locus plays a major role to inhibit phagosome maturation.

**Fig. 7:**
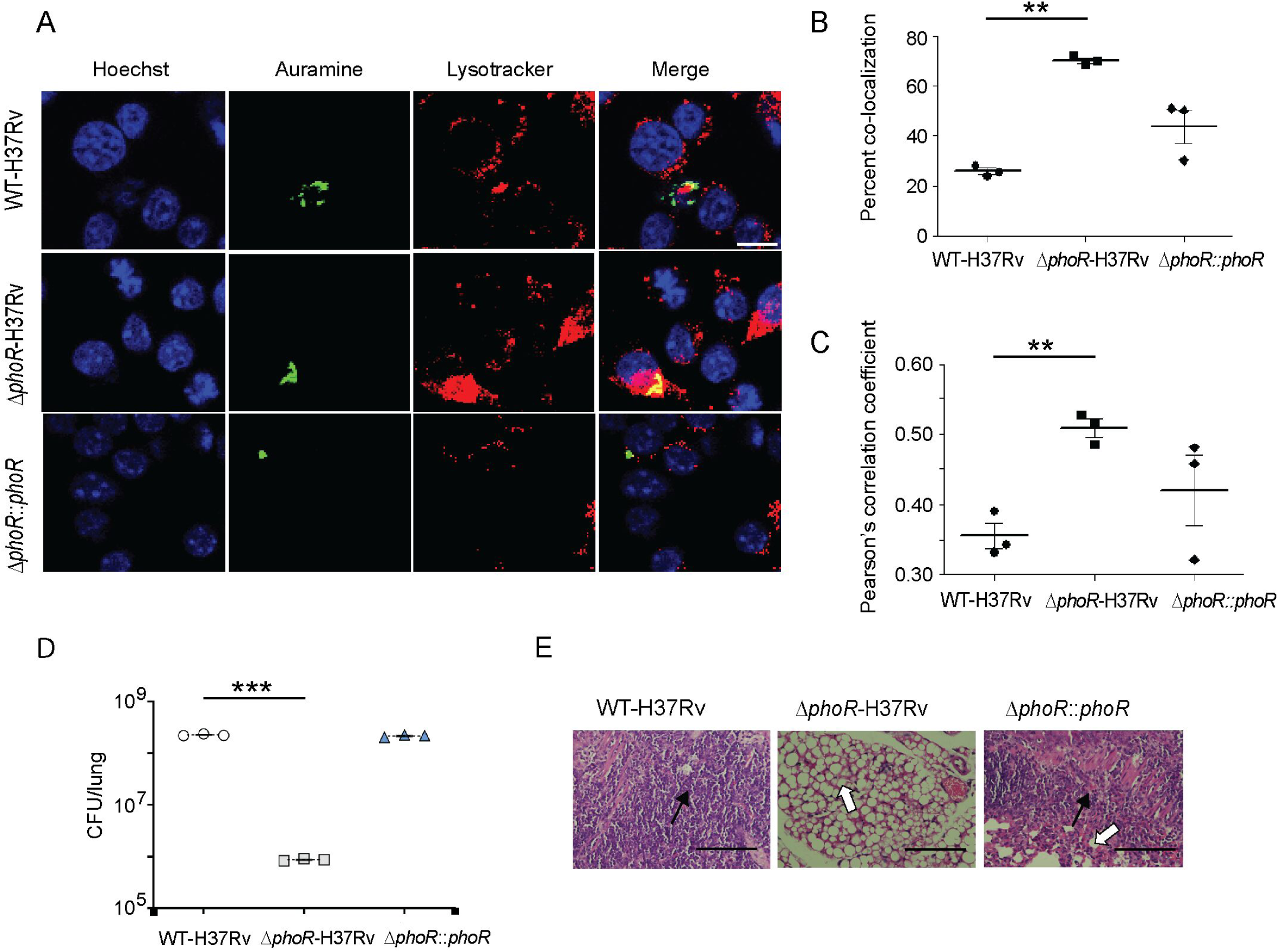
PhoR contributes to mycobacterial survival in cellular and animal models. (A) WT-H37Rv and *ΔphoR*-H37Rv were used to infect murine macrophages. Mycobacteria and host cells were stained with phenolic auramine solution, and LysoTracker respectively. Two fluorescence signals (Mycobacterial strains: green; LysoTracker: red) and their merging are displayed by confocal images (scale bar: 10 µm). (B) The % colocalization of auramine labelled strains with Lysotracker was measured in 10 different fields by counting at least 100 infected cells (***P≤ 0.001). (C) The data present Pearson’s correlation coefficient of images displaying internalized auramine-labelled mycobacteria and Lysotracker red marker in macrophages. Average values with standard deviations were obtained from three independent experiments (*P<0.05; ***P<0.001). (D) Mycobacterial survival *in vivo* was measured in mice 4 weeks post-infection with ~100 CFU / lung of bacterial load. CFU in the lungs represent average values with standard deviations from three to five animals for each bacterial strains (***P≤0 0.001). (E) For histopathology after 4 weeks of infection with indicated mycobacterial strains, lung sections were stained with hematoxylin and eosin, observed under a light microscope, and images of the sections collected at x40 magnification display granulomas (filled arrows) and alveolar space (empty arrows) (scale bar: 200 µm).

The reduced ability of *ΔphoR*-H37Rv to survive in macrophages led us to investigate its ability to survive *in vivo*. To examine this, mice were infected with WT-H37Rv and *ΔphoR*-H37Rv. CFU analyses after day 1 of infection revealed presence of comparable counts of both WT-H37Rv and the *ΔphoR*-H37Rv bacilli (~100 bacilli) in the mice lungs. However, after 4 weeks of infection, the PhoR-deleted mutant showed a ~25-fold lower bacterial burden in mice lung relative to WT-H37Rv (Fig. 7D). The fact that *phoR* locus contributes to pathogenesis was further confirmed as *ΔphoR*-H37Rv expressing a stable copy of the *phoR* gene showed a lung burden comparable to that of WT-H37Rv (1.05 ±0.1-fold difference). From these results we conclude a significantly reduced ability of *ΔphoR*-H37Rv to replicate in mice. Also, histopathological evaluations suggest that PhoR-deleted H37Rv shows a significantly less severe pathology relative to WT-H37Rv (Fig. 7E). In conjunction with the previous study (Xing *et al*., 2017), these results unambiguously suggest that *M. tuberculosis pho*R locus remains essential for mycobacterial pathogenesis.

## Discussion

The first suggestion that *M. tuberculosis phoPR* system might be involved in pH sensing was made by Smith and co-workers, when they had reported PhoP-dependent regulation of numerous pH-sensitive genes in the transcriptome analyses (Walters *et al*., 2006). Later, Russell and co-workers have shown a significant overlap between phagosomal acidic pH regulon and *phoPR* regulon (Rohde *et al*., 2007). These results led to the identification of a unique *M. tuberculosis*-specific acid and phagosome regulated *aprABC* locus (Abramovitch *et al*., 2011). Importantly, *aprABC* shows a *phoPR*-dependent activation during mycobacterial growth under acidic pH and contributes to remodel the phagosomal environment for effective intracellular adaptation of the bacilli (Abramovitch *et al*., 2011). The other suggested possibility was that *phoPR* responds to a non-conventional intra-bacterial signal associated with altered physiology at acidic pH, such as change in carbon metabolism and/or redox homeostasis (Baker *et al*, 2019; Baker *et al*., 2014). Although we now know the *phoPR* regulon is induced under acidic pH *in vitro* and in macrophages (Abramovitch *et al*., 2011; Bansal *et al*., 2017), what remained unknown is whether PhoPR directly responds to low pH conditions.

While our present knowledge on the regulatory stimuli for a few SKs is often derived from transcriptomic data, for majority of SKs the environmental signals that directly regulate their activity remain unknown. This lack of knowledge accounts for our limited understanding on the contribution of these systems to different host niches in the context of mycobacterial pathogenesis. In this study, we observed a significantly elevated level of phosphorylation of PhoR under acidic pH, and high salt conditions of growth relative to normal conditions (Fig. 1A). Importantly, under identical stress conditions we observed a significantly elevated phosphorylation of PhoP (Fig. 1B), suggesting a stress-specific likely phospho-transfer from PhoR to PhoP. Further, we demonstrated that P~PhoP is almost undetectable in *ΔphoR*-H37Rv even under acidic pH (Fig. 2), allowing us to conclude that acidic pH/high salt condition is directly sensed by PhoR, and low pH promotes PhoR-dependent PhoP phosphorylation. Consistent with these results, transcriptome analyses of *ΔphoR*-H37Rv and WT-H37Rv demonstrate a strikingly differential regulation of low pH-inducible PhoP regulon with a significant impact of PhoR on *in vivo* recruitment of PhoP within acidic-pH inducible promoters (Fig. 3). Collectively, these results suggest that lack of acidic pH-inducible expression of PhoP regulon in *M. tuberculosis* is attributable to the absence of PhoR.

Numerous studies were undertaken to investigate cross-talk between mycobacterial TCS proteins (Agrawal *et al*., 2015; Lee *et al*., 2012; Malhotra *et al*., 2015). Although these studies suggest “multi-to-one” signalling relationships, the physiological consequences of these interactions to environmental adaptations, or how they facilitate important cellular functions remain unknown. In recent years a number of regulators have been shown to interact with PhoP impacting mycobacterial physiology (Anil Kumar *et al*., 2016; Bansal *et al*., 2017; Sevalkar *et al*., 2019; Singh *et al*, 2020b).However, till date phosphorylation of PhoP with a non-cognate kinase have not been documented. As we begin to understand the contribution of TCS proteins to mycobacterial physiology beyond the classical phospho-transfer pathway, it becomes evident that these systems participate in extensive signalling networks. These pathways may have important consequences on pathogenesis particularly in view of the fact that many of the environmental conditions encountered by *M. tuberculosis* are part of the same niche, suggesting that cross-regulation between concomitantly activated TCSs may be one of the common mechanisms pathogenic mycobacteria employ to facilitate environmental adaptation. To investigate whether PhoP is activated by any other SK(s), using M-PFC experiments (Fig. 4) coupled with phosphorylation assays (Fig. 5), we demonstrate specific cross-talk of PhoP with PrrB. Although these results appear to be in keeping with the role of *phoP* locus in mycobacterial nitrogen metabolism (Singh *et al*., 2020b), we were unable to detect *in vivo* phosphorylation of PrrB (Fig. 5E) under nitrogen limiting conditions. However, we unexpectedly observed PrrB undergoing partial phosphorylation *in vivo* under acidic pH conditions of growth relative to normal pH (compare lanes 1 and 3, Fig. 5E). Together, this counterintuitive finding strikingly accounts for PhoR-independent, but PhoP-mediated regulation of a few pH-inducible genes (Fig. 3A; Figs. 5C-D and Fig. S3B). Note that the environmental signal(s) that triggers specific and non-canonical mode of PhoP activation via PrrB, requiring an appropriate transcriptional response, remains unknown. We speculate that *prrB* responds to a non-conventional signal associated with altered physiology at acidic pH. Nevertheless, these findings connect two SKs to a single RR (PhoP) in a stress-specific manner and provide new biological insights into mechanism of activation of the versatile regulator.

Together, these results uncover that a SK from one TCS communicates with a non-cognate RR of another TCS to expand the signalling repertoire of mycobacteria. In this “multi-to-one” signalling pathway, promiscuous SKs phosphorylate more than one RRs, impact input recognition capabilities of RR and subsequently expand and/or fine tune downstream response by the phosphorylated RR. We propose that the multi-stimulus SK-dependent regulation of a RR likely reflects the need of the pathogen to fine-tune its complex physiology within the colonized niche. It is possible that rather than an isolated exception, RRs getting activated in response to multiple signals is not so uncommon. In fact, fewer TCSs of *M. tuberculosis* relative to other bacterial species of comparable genome size is suggestive to be part of a more efficient regulatory system. With more insights appreciating host adaptation of the tubercle bacilli which rely on bacterial TCSs - is an area of growing interest that holds strong potential to develop adjunct therapy. It is noteworthy that ethoxzolamide, which inhibits PhoPR virulence associated regulon in *M. tuberculosis* (Johnson *et al*, 2015), has been shown to reduce bacterial burden in both infected macrophages and mice. Although the mechanism is yet to be understood, ethoxzolamide is thought to inhibit PhoR sensing by targeting cell surface carbonic anhydrases.

To investigate regulation of P~PhoP in mycobacteria, we next discovered that PhoR functions as a robust phosphatase of P-PhoP, suggesting that the sensor in addition to its kinase function undertakes an additional layer of regulatory function to inhibit ‘triggering ON’ of the PhoP regulon unless it is necessary. The finding that PhoR is a dual function sensor protein with both kinase and phosphatase functions (Figs. 5-6) provides an explanation to how PhoP regulon remains repressed under non-inducing conditions. We consider two major physiological significances of the newly-identified function of PhoR. First, it remains an effective mechanism to prevent PhoP regulon expression which perhaps is only needed during mycobacterial adaptation to low pH. Secondly, restoring PhoP to its unphosphorylated status by PhoR may be essential to reverse pH regulon under normal conditions that support increased metabolic activity. It is noteworthy that PhoP plays a major role in integrating low pH conditions and redox homeostasis by controlling mycobacterial metabolic plasticity (Baker *et al*., 2019; Baker *et al*., 2014). Thus, PhoR controls the net phosphorylation status of PhoP and determines the final output on mycobacterial adaptive programme to low pH conditions, most likely through its contrasting kinase and phosphatase activities, facilitates an integrated view of our results. The balancing act of two activities, which controls the expression of acidic pH-inducible PhoP regulon is schematically shown as a model in Fig. 6J. The result identifying E260, adjacent to the phosphorylation site H259, as a critically important residue for PhoR phosphatase function (Fig. 6H), provides us with a fundamental biological insight into how a limited number of residues of the DHp domain contribute to two contrasting functions of the sensor protein and determine environment-specific net activation status of the cognate RR.

In conclusion, our results for the first time demonstrate that acidic pH activates PhoP by promoting its phosphorylation in a PhoR-dependent manner, and therefore, expression of PhoP regulon requires the presence of PhoR. We also identify a non-canonical mechanism of activation of PhoP, connecting two SKs with signal dependent activation of the virulence regulator. Our results further uncover phosphatase activity of PhoR, and identify the motif and residues responsible for kinase/phosphatase (dual) functioning of the SK. Collectively, we conclude that both kinase and phosphatase functions of PhoR remain critically important for downstream functioning of the activated regulator, and propose a model suggesting how dual functioning of PhoR effectively regulates final output of the PhoP regulon in an environment-dependent manner. In keeping with these results, PhoR remains essential for mycobacterial virulence in both cellular and animal models (Fig. 7), and together, these results have striking implications on the mechanisms of virulence regulation.

## Materials and Methods

### Bacterial culture, and growth experiments

*E. coli* DH5α, and *E. coli* BL21 (DE3) were utilized to clone and express mycobacterial proteins, respectively. Construction of PhoR-deleted *M. tuberculosis* (*ΔphoR*-H37Rv) using ‘mycobacterial recombineering’ and complementation of the mutant have been described in this study. PhoPR-deleted *M. tuberculosis* (*ΔphoP*-H37Rv) and the complemented mutant have been described previously (Walters *et al*., 2006). *M. tuberculosis* strains were grown aerobically in 7H9 liquid broth or on 7H10-agar medium as described (Goyal *et al*., 2011). Wild-type (WT) or mutant *M. tuberculosis* strains were transformed and selected in presence of antibiotics as described (Jacobs *et al*, 1991). For stress conditions due to low pH and salt, cells were grown to OD_600_ of 0.4, washed with 7H9 buffered with 2N HCl for pH 7.0 or pH 4.5, or 7H9 containing 250 mM sodium chloride, respectively, re-suspended in media of indicated pH and salt, and grown for additional 2 hours at 37°C. For redox stress, mycobacterial cells were inoculated into 7H9-ADS (albumin-dextrose-sodium chloride) containing ~5 mM diamide (Sigma) at OD_600_ of 0.05, and grown for additional 48 hours at 37°C as described (Goar *et al*, 2022). For mycobacterial growth under nitrogen limiting conditions, cells were grown in synthetic 7H9 medium using 0.04 g/L ferric citrate and 0.5 g/L sodium sulphate instead of comparable concentrations of ferric ammonium sulphate, ammonium sulphate and 0.5 g/L glutamic acid, as nitrogen sources.

### Cloning

His-tagged PhoP was cloned between BamHI and PstI sites of both p19Kpro ((De Smet *et al*., 1999) and pSTKi (Parikh *et al*, 2013) using the same primer pair FPmphoP/RPmphoP (Table S1). Likewise, full-length PhoR, and PrrB were cloned in pSTKi and p19Kpro between NdeI/PstI and BamHI/HindIII sites, using primer pairs FPphoRcom/RPphoRcom and FPprrB/RPprrB, respectively. *M. tuberculosis* PhoP, PhoR, and PrrB proteins carrying an N-terminal His_6_-tag were expressed in mycobacteria as described previously (Anil Kumar *et al*., 2016). Cloning of cytoplasmic domain of PhoR (PhoRC comprising residues 193-485 of PhoR), full-length DosR, and PhoP and expression in *E. coli* have been described earlier (Gupta *et al*., 2006; Singh *et al*., 2020a). To express His_6_-tagged cytoplasmic domain of PrrB (PrrBC, comprising residues 200-446 of PrrB), the amplicon was inserted in pET28b between NdeI and HindIII sites, and PrrBC was expressed as described for PhoRC (Gupta *et al*., 2006). Likewise, NdeI and HindIII sites of pET28b (Novagen) were used to construct plasmid expressing PrrA. Specific point mutations within the indicated ORFs were introduced by overlap extension PCR, and constructs were checked by DNA sequencing. The sequences of the oligonucleotides used for cloning and the plasmids used for expression are listed in Tables S1, and S2, respectively.

### Expression and purification of Proteins

Plasmids expressing recombinant WT or mutant PhoP, DosR, PrrA, cytoplasmic domains of PhoR and PrrB proteins, were expressed as fusion proteins containing N-terminal His_6_-tag and purified by Ni-NTA chromatography as described (Gupta *et al*, 2009; Gupta *et al*., 2006). In each case, protein purity was examined by SDS-PAGE, and concentration was assessed by Bradford reagent (with BSA as a standard). The storage buffer for proteins comprised of 50 mM Tris-HCl, pH 7.5, 300 mM NaCl, and 10% glycerol.

### *In vitro* phosphorylation of proteins

For autophosphorylation, WT or mutant PhoRC, and PrrBC proteins (~ 3 µM) were included in a phosphorylation mix (50 mM HEPES, pH 7.5, 50 mM KCl, 10 mM MnCl_2_) which comprised of 25 μM of (γ-^32^P) ATP (BRIT, India). The reactions were allowed to continue for 1 hr at 25°C. To initiate phosphotransfer, ~ 3 µM PhoP or its truncated variants were included in the phosphorylation, the reactions continued at 18°C for indicated times, and terminated by 10 mM EDTA. The reactions products were analysed by SDS/polyacrylamide gels, gels were dried, and signals digitized by a phosphorimager.

### *In vivo* protein phosphorylation experiments

Cell lysates were resolved in polyacrylamide gels containing acrylamide–Phos-tag ligand (Wako Laboratory, Japan) as per manufacturer’s recommendation to detect PhoP/PhoR proteins and their phosphorylated forms. These gels were copolymerized in presence of 50 µM Phos-tag acrylamide and 100 mM MnCl_2_. *M. tuberculosis* cell-lysates were prepared as described previously (Anil Kumar *et al*., 2016) and total protein content determined by Bradford reagent. Standard running buffer [0.4% (w/v) SDS, 25 mM tris, 192 mM glycine] was used for electrophoresis of samples on Phos-tag gels for 5 hours under 20 mA. Next, the resolved samples were transferred to PVDF membrane and detected by Western blot analyses. Anti-His antibody was from GE Healthcare; anti-PhoP antibody was elicited in rabbit (AlphaOmegaSciences); goat anti-rabbit and goat anti-mouse secondary antibodies were from Abexome Biosciences, and the chemiluminescence reagent to develop the blots was from Millipore.

### Construction of *ΔphoR-*H37Rv

PhoR-deleted *M. tuberculosis* H37Rv was constructed by ‘mycobacterial recombineering’ (van Kessel & Hatfull, 2007). To construct *phoR* allelic exchange substrate (AES), left homology region (LHR) encompassing −970 to +30 (relative to *phoR* ORF) and the right homology region (RHR) encompassing +1423 to +2423, were PCR amplified using the primer pairs FPphoRLHR/ RPphoRLHR and FPphoRRHR/ RPphoRRHR, respectively. The primers included PflM1(LHR) and DraIII (RHR) restriction sites, and digestion with these enzymes results in ends that are compatible for cloning with Hyg^r^ cassette (derived from modified-pENTR/D-TOPO plasmid) and OriE+ cosλ fragment (derived from p0004S plasmid). Also, ScaI site was inserted both in the LHR forward primer and RHR reverse primer. Next, amplicons of LHR and RHR were digested with PflM1 and DraIII, respectively; and the resultant fragments were ligated with Hyg^r^ cassette (1.3 kb) and OriE+ cosλ fragment (1.6 kb) to generate *phoR* AES. Next, it was digested with ScaI to generate linear *phoR* AES (~3.3 kb) suitable for ‘recombineering’. Fig. 2A shows the schematic map of *phoR* AES plasmid.

For homologous recombination, WT-H37Rv strain was electroporated with pNit-ET (a kind gift of Prof. Eric Rubin) expressing gp60 and gp61 to generate recombineering proficient H37Rv::pNit-ET strain. This strain was grown to OD_600_ ≍0.4 and 0.5 µM isovaleronitrile (Sigma) induced expression of the recombinases. Next, 200 ng of *phoR* linear AES was used to electroporate cells and transformed cells were grown on to 7H10/OADC agar plates containing hygromycin. Genomic DNAs were isolated from single colonies and screened for *phoR* replacement. To construct the complemented mutant, *phoR* encoding ORF was PCR amplified using the primer pair FPphoRcomp/RPphoRcomp, digested with NdeI/PstI and ligated to double digested pST-Ki (Parikh *et al*., 2013). Next, pST-*phoR* clone was confirmed by sequencing, electro-transformed in *ΔphoR*-H37Rv competent cells, and the transformed colonies were selected on 7H10/OADC/Kan/Hyg. The mutant and the complemented strains were verified by gene-specific PCR amplification using corresponding genomic DNAs as PCR templates. We also performed Southern blot hybridization to confirm the mutant, and RT-qPCR experiments compared *phoR* mRNA levels of WT-H37Rv, mutant, and the complemented strain.

### Southern blot hybridization

This approach was used to confirm *ΔphoR*-H37Rv mutant. Approximately 1 µg of genomic DNA of WT and *ΔphoR*-H37Rv were restricted using BamHI and separated by agarose gel electrophoresis. Next, the DNA was transferred onto the Immobilon membrane (Millipore) by vacuum-based blotting, and hybridized with radio-labelled gene-specific probes. The *hrcA* and *phoR*-specific probes (Table S1) were generated by PCR using alpha ^32^P-dCTP (BRIT, India) and oligonucleotide primers, which were used to clone the respective ORFs. Pre-hybridization, hybridization and washing steps were in accordance with the standard protocol. The image was developed and digitized with a phosphorimager (GE Health care).

### Construction of *phoR* and *prrB* knock-down mutants of *M. tuberculosis* H37Rv

In this study, we utilized a previously-described CRISPRi-based strategy (Singh *et al*., 2016) to construct knock-down mutants of *phoR* and *prrB*. This approach inhibits expression of genes via inducible expression of dCas9 protein using target gene-specific guide RNAs (sgRNA). First, WT-H37Rv was transformed with *S. pyogenes* dCas9 expressing integrative plasmid pRH2502 to generate *WT-H37Rv::dCas9*. Next, 20-nt long *phoR* and *prrB*-targeting spacer sgRNAs were cloned in pRH2521 using BbsI enzyme and the constructs sequenced. The oligonucleotides were designed such that the expressed sgRNA comprises of a 20 bp sequence that remains complementary to the non-template strand of the target gene. Finally, the corresponding clones were electroporated into *M. tuberculosis* harbouring pRH2502. To express dcas9 and repress sgRNA-targeted genes (*phoR* or *prrB*), the bacterial cultures were grown in Middlebrook 7H9 broth, supplemented with 0.2 % glycerol, 10 % OADC, 0.05 % Tween, 50 μg/ml hygromycin and 20 μg/ml kanamycin at 37°C, anhydrotetracycline (ATc) was added every 48 hours to a final concentration of 600 ng/ml, and cultures were continued to grow for 4 days. Next, RNA isolation was carried out, and RT-qPCR experiments verified repression of target genes. For the induced strains (in the presence of ATc) expressing sgRNAs targeting +234 to +253 (relative to *phoR* translational start site) and +56 to +75 sequences (relative to *prrB* translational start site), we obtained approximately 92% and 95% reduction of *phoR* and *prrB* RNA abundance, respectively, compared to the respective un-induced strains. The oligonucleotides used to generate gene-specific sgRNA constructs and the plasmids utilized in knock-down experiments are shown in supplementary Tables S1 and S2, respectively.

### RNA isolation

Isolation and purification of RNA from *M. tuberculosis* strains have been described previously (Khan *et al*, 2022). To ensure removal of genomic DNA, each RNA preparation was incubated with RNase-free DNaseI at room temperature for 20 minutes. RNA concentrations were measured by recording absorbance at 260 nm, and sample integrity was verified using formaldehyde-agarose gel electrophoresis by assessing intactness of 23S and 16S rRNA.

### RNA sequencing and data analysis

RNA-sequencing and data analysis have been performed by AgriGenome (India). Agilent 2200 system examined RNA integrity, and ‘si-tools Pan-prokaryote ribopool probes removed rRNA. TruSeq standard RNA library prep kit was used to prepare the libraries, which were sequenced by Illumina HiSeq x10 platform to generate 150-bp paired-end reads. Details of data analyses was as described (Goar *et al*., 2022). Fold changes ≥2 or ≤-2 of treated (compared to control) samples were used to generate heat-maps using GENE-E software. Our results showing a comprehensive list of genes have been submitted in the NCBI’s database (GEO accession number GSE180161).

### Quantitative Real-Time PCR

Total RNA extracted from *M. tuberculosis* cultures grown under specific conditions were used for cDNA synthesis and PCR reactions using Superscript III platinum-SYBR green one-step RT-qPCR kit (Invitrogen). Details of PCR conditions are described previously (Goar *et al*., 2022), and *M. tuberculosis* RpoB or 16S rDNA was used as endogenous controls. Two independent RNA preparations were always used to evaluate fold difference in gene expression using ΔΔC_T_ method (Schmittgen & Livak, 2008). Table S7 lists sequences of oligonucleotide primers, which were utilized to determine specific mRNA levels relative to control set displaying no differential expression. Enrichment attributable to PhoP binding to its target promoters were assessed by using 1 µl of IP or mock IP (no antibody control) DNA with SYBR green mix (Invitrogen) and promoter-specific primers.

### ChIP-qPCR

Details of ChIP-qPCR experiments using anti-PhoP antibody has been described elsewhere (Khan *et al*., 2022). *In vivo* recruitment of the regulator was assessed using IP DNA in a reaction buffer containing SYBR green mix (Invitrogen), appropriate PAGE-purified primers and one unit of Platinum Taq DNA polymerase (Invitrogen). Amplifications were done for 40 cycles using Applied Biosystems real-time PCR detection system, and signal from an IP without antibody was compared to measure efficiency of recruitment. PCR-enrichment specificity from identical IP samples was also verified using 16S rDNA or RpoB-specific primers. A single product was amplified in all cases, and duplicate measurements were made in each case from two independent bacterial cultures.

### Mycobaterial protein fragment complementation (M-PFC) assays

*M. tuberculosis phoP* was expressed from pUAB400 (Kan^R^), and transformed cells (*M. smegmatis* mc^2^155 containing pUAB400-*phoP*) were grown in liquid medium to obtain competent cells. Likewise, sensor kinases were expressed from the episomal plasmid pUAB300 (Hyg^R^) after cloning their ORFs between BamHI/HindIII sites. Sequences of the oligonucleotide primers and relevant plasmids are listed in Table S8 and Table S9, respectively. Each construct was checked by DNA sequencing, and M. smegmatis cells carrying both plasmids were selected on 7H10/Kan/Hyg plates in presence or absence of 15 µg/ml Trimethoprim (TRIM) as described earlier (Singh *et al*., 2006). *phoP*/*phoR* co-expressing constructs were used as a positive control (Singh *et al*, 2014).

### Macrophage Infections

RAW264.7 macrophages were infected with titrated cultures of H37Rv, *ΔphoR*-H37Rv and complemented mutant strain at a multiplicity of infection (MOI) of 1:5 or 1:10 for as described previously (Anil Kumar *et al*., 2016). While phenolic auramine solution was used to stain *M. tuberculosis* H37Rv strains, the cells were stained with 150 nM Lyso-Tracker Red DND 99 (Invitrogen). Next, cells were fixed, analysed using confocal microscope (Nikon, A1R), and processing of digital images were carried out with Nikon image-processing software. Details of the experimental methods and the laser/detector settings were optimized using macrophage cells infected with WT-H37Rv as described (Anil Kumar *et al*., 2016). A standard set of intensity threshold was made applicable for all images, and percent bacterial co-localization was determined from three biological repeats by analyses of more than 100 bacteria per sample from at least five random fields.

### Mouse infections

Animal handling protocols were approved by the Institutional Animal Ethics Committee (IAEC/17/05, and IAEC/19/02). Independently passaged *M. tuberculosis* H37Rv cultures from mid log phase were used to infect C57BL/6 mice (8-10 weeks old) by inhalation-based Glass-col exposure system. The animals were sacrificed by cervical dislocation, lungs isolated aseptically from the euthanized animals were homogenized in sterile 1X PBS and plated on 7H11 agar plates after serially diluting the lysates. The plates contained 10% OADC and following antibiotics: 50 μg/ml carbenicillin, 30 μg/ml polymyxin B, 10 μg/ml vancomycin, 20 μg/ml cycloheximide, 20 μg/ml Trimethoprim, and 20 μg/ml Amphotericin B. The bacterial load was determined 1-day post infection from the infected mice, and for each strain deposition of 100-200 CFU of bacilli was found in the lungs. Finally, mice were euthanized 4-weeks post infection, lungs and spleens were plated on 7H11 plates supplemented with a 0.5% glycerol, and 10% albumin-dextrose-catalase (ADC, Middlebrook) to determine CFU. After fixing left lung lobes in 10% buffered formalin, these were embedded in paraffin and stained with haematoxylin and eosin for visualization under the microscope, and percent pathology of the infected lungs were calculated using Image J software.

## ACKNOWLEDGMENTS

We acknowledge G. Marcela Rodriguez and Issar Smith (PHRI, New Jersey Medical School - UMDNJ) for Δ*phoP*-H37Rv, and the complemented mutant strain. We thank Divya Arora and Vinay Nandicoori from National Institute of Immunology (NII) and Ritesh Sevalkar from our laboratory for their help in constructing *ΔphoR*-H37Rv. We acknowledge Rajni Garg, and Anunay Sinha from the laboratory of Sanjeev Khosla for their help in constructing the knock-down mutants, and analysis of RNA sequencing data, respectively. We thank Neeraj Khatri and members of the Institutional animal facility for their assistance to obtain clearance of our projects from the Animal Ethics Committee. This study received financial support from CSIR-IMTECH intramural grant OLP-0170, CSIR-funded FBR project MLP-0049, and SERB-funded project (EMR/2016/004904) to D.S. P.R.S., was supported by SERB (DST); H.G., P.P., K.M., B.B., A.K.V., and H.K. were supported by CSIR; R. B. was supported by UGC. The funders were not part of study design, collection of data, interpretation of results, or our decision to submit the work for publication.

## Author contributions

P.R.S., H.G., P.P., K.M., B. B., A.K.V., R. B. and D.S. designed research; P.R.S., H.G., P. P., K.M., B. B., A.K.V., R. B., and H. K., performed research; S.K. contributed analytical tools; P.R.S., H.G., P. P., K. M., B. B., A.K.V., R. B., H. K., and D.S. analysed data; and D.S. wrote the manuscript

## Conflict of Interest

The authors declare no conflict of interest.

## Data availability

All RNA sequencing data have been deposited in the GEO database with accession number GSE180161. All other relevant data are part of the main text and its supporting Information files.

